# Hierarchical Processing of Natural Scenes in the Human Pulvinar

**DOI:** 10.1101/2025.03.20.644381

**Authors:** Daniel R. Guest, Emily J. Allen, Kendrick N. Kay, Michael J. Arcaro

**Author notes:** Co-senior author (Kendrick N. Kay, Michael J. Arcaro).

## Abstract

The hierarchical organization of the ventral visual cortex has been the focus of theories and computational models characterizing high-level visual processing and object recognition, often overlooking potential contributions of subcortical structures. The pulvinar, through its extensive reciprocal connections with ventral visual cortex, is well-positioned to play a prominent role in high-level visual recognition processes. Here, we investigated whether the pulvinar plays such a role using a high-resolution 7T fMRI dataset of responses to tens of thousands of natural scenes. Encoding models testing the representation of different stimulus features revealed a pulvinar region selective for bodies and faces presented within the contralateral visual hemifield. Model-free analyses demonstrated that this region is predominantly co-active with body- and face-selective cortical areas during natural scene viewing. This functional specificity was embedded within a broader gradient of cortical correlations across the pulvinar mirroring the hierarchical organization of ventral visual cortex. These findings challenge cortico-centric models of object vision and implicate a role of the pulvinar in high-level vision. More broadly, these results demonstrate that principles of cortical organization, including functional clustering and hierarchical organization, also manifest in subcortex, and highlight the value of using naturalistic stimuli to probe visual function.

## Introduction

Higher visual functions, particularly object and face recognition, are crucial for interacting with and making sense of the environment. A hallmark of primate vision is our ability to rapidly and effortlessly identify and categorize objects across variations in the retinal input (DiCarlo et al., 2012). This remarkable capacity relies on the brain’s ability to extract relevant visual features from complex visual input and integrate them into a coherent percept.

The hierarchical organization of the ventral visual cortex has long been recognized as a key factor in transforming retinal input into useful perceptual representations (DiCarlo & Cox, 2007). This hierarchy progresses from early visual cortex (V1, V2) responding to low-level features such as local contrast and orientation, through intermediate areas (V4) encoding more complex features such as textures and shapes, to anterior regions exhibiting categorical distinctions in their responsiveness to faces, bodies, and objects (Grill-Spector & Weiner, 2014; Kourtzi & Connor, 2011; Roe et al., 2012). In addition to feedforward pathways giving rise to increasingly complex representations, recurrent and feedback connections are thought to help ensure robust recognition in the face of challenges during natural visual experience, such as partial occlusion or varying light conditions (Kar & DiCarlo, 2021; Kay & Yeatman, 2017).

However, this cortico-centric view is incomplete. Recent evidence suggests that recurrent connectivity with the thalamus, particularly the pulvinar, may play a critical role in achieving robust perceptual and cognitive functions (Halassa & Kastner, 2017; Reinhold et al., 2015; Sherman, 2017). The pulvinar, with its extensive reciprocal connections to visual cortex, is well-positioned to influence visual processing at multiple levels. While much research has focused on the pulvinar’s involvement in early sensory processing (Bender, 1981; Petersen et al., 1985; Purushothaman et al., 2012), visual attention (Kastner et al., 2004; Saalmann et al., 2012), and visuomotor behaviors (Mundinano et al., 2018; Wilke et al., 2010), its distinct anatomical connectivity with ventral visual cortex (Webster et al., 1993) suggests a potential role in high-level object recognition processes.

Despite the presence of distinct anatomical connectivity between the pulvinar and cortical regions supporting high-level visual processes, our understanding of the pulvinar’s role in complex visual functions, such as object recognition, remains limited. Previous investigations of pulvinar function have predominantly employed simple stimuli, focusing on low-level visual features, such as orientation and image contrast (Bender, 1981; Gattass et al., 1978). Some evidence hints at the pulvinar’s involvement in higher-level visual representations: anatomical tracing in non-human primates reveals connections between the pulvinar and face-selective cortical regions (Grimaldi et al., 2016); electrophysiological recordings have identified pulvinar neurons sensitive to face-like shapes (Nguyen et al., 2012) and facial emotion (Maior et al., 2010); and fMRI studies in humans have identified functional substructure within the pulvinar with sensitivity to visual categories (Arcaro et al., 2018; Wen et al., 2023). However, the specificity of these findings to object recognition processes remains unclear. For example, medial pulvinar damage can impair emotion recognition, while sparing other aspects of visual recognition (Ward et al., 2007). Moreover, individual pulvinar regions receive convergent inputs from functionally diverse cortical regions, potentially resulting in broadly tuned representations. Thus, how the pulvinar represents complex visual information, its functional topography, and its relationship to cortical processing of visually presented objects remains unclear.

To investigate the pulvinar’s role in object recognition and the relationship of the pulvinar to cortical areas, we leveraged the Natural Scenes Dataset (NSD; http://naturalscenesdataset.org), a 7T fMRI dataset consisting of responses to tens of thousands of natural scenes. NSD constitutes an especially promising opportunity to study the pulvinar, given the use of ultra-high magnetic field strength (7T) to improve signal-to-noise ratio and achieve high spatial resolution (1.8 mm). We first developed encoding models to characterize spatial coding and responsiveness to different visual features present within natural scenes, including low-level image content and selectivity to specific visual categories. We then used model-free correlation analyses to identify co-activation patterns between the pulvinar and visual cortex during the processing of natural scenes. These analyses revealed a pulvinar region selectively responsive to faces and bodies, distinct from portions of the pulvinar sensitive to low-level visual content. Notably, both regions exhibited retinotopic organization, suggesting a common organizing principle across different levels of visual processing. The body- and face-selective pulvinar regions also showed specific co-activation patterns with corresponding body- and face-selective cortical areas. Furthermore, these distinct functional clusters were embedded within a broader gradient of cortical correlations across the pulvinar, mirroring the hierarchical organization of ventral visual cortex. These findings position the pulvinar as an important structure in high-level vision, potentially interfacing with cortex at all levels of visual processing.

## Results

### Using pRF models to characterize representations of features in natural scenes

We first characterized the pulvinar’s response to different visual features within natural scenes by building and testing encoding models for the Natural Scenes Dataset (NSD). We evaluated a wide range of image features, including low-level properties, such as local contrast, and high-level content, such as bodies and faces. To predict voxel-wise responses, we coupled features extracted from the NSD images with a population receptive field (pRF) model (Kay, Winawer, et al., 2013) (Figure 1). This approach characterizes each voxel as being jointly tuned for a specific stimulus feature and an area of visual space. We tested models corresponding to seven different features prevalent in real-world scenes: local contrast, salience, faces, bodies, words, foreground, and background (Supplementary Figure 1; see Methods for details about how each feature was computed). For each feature, we obtained a full set of best-fitting pRF parameters, including pRF position, size, and gain for every voxel within each participant. This model-based approach allowed us to assess multiple levels of visual processing within the pulvinar.

**Figure 1.**
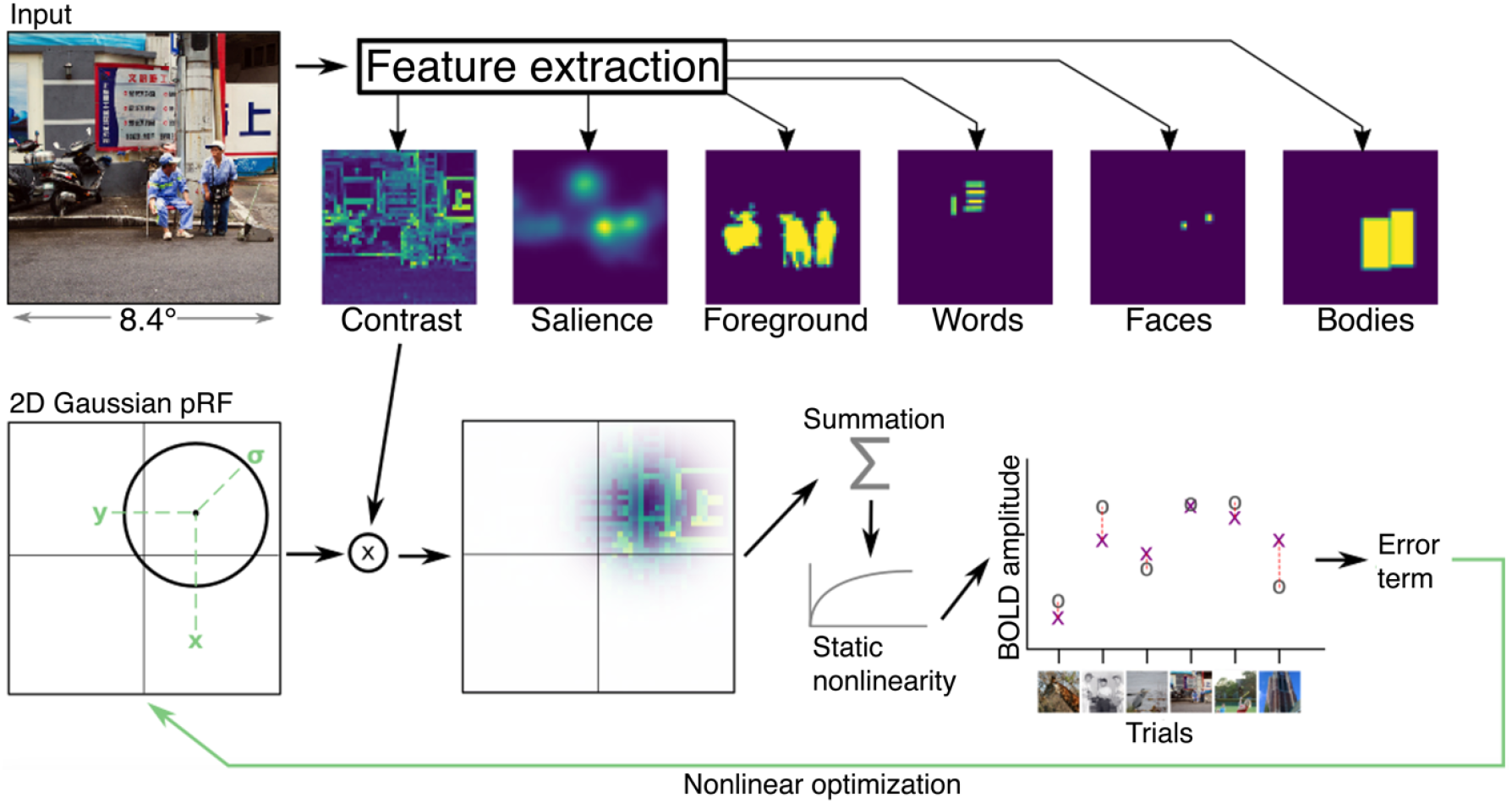
Identifying stimulus features encoded in voxel responses using population receptive field (pRF) models. Each NSD image was converted into feature maps that indicated the presence of a given feature at each location in the image. Seven types of feature maps, corresponding to different low-, mid-, and high-level image properties (see Methods for details), were prepared (background, not shown, was defined as the inverse of foreground). To predict the response to an image, a given feature map is weighted by a 2D isotropic Gaussian, summed over the visual field, and then a compressive static nonlinearity is applied (Kay, Winawer, et al., 2013). Model parameters are optimized to minimize error between observed BOLD amplitudes (gray circle markers) and predicted BOLD amplitudes (purple “X” markers). For each voxel, a separate model was fit for each feature type, attempting to account for the fMRI response amplitudes to the 9,000–10,000 natural scenes seen by each NSD subject.

### Distinct pulvinar regions responsive to low- and high-level features

Our analyses revealed distinct regions within the pulvinar responsive to different types of visual content within natural scenes. We observed a clear functional segregation between regions of the pulvinar that process low-level visual features and those responsive to high-level object categories (Figure 2). While the variance explained was relatively low in the pulvinar compared to typical cortical BOLD responses, this is likely attributable to the high levels of physiological and thermal noise inherent in midbrain structures (Barry et al., 2013). Despite this, the observed patterns of activation were consistent across individuals, underscoring the reliability and functional relevance of these distinctions. Across subjects, low-level feature processing was predominantly localized to the inferior-lateral portions of the pulvinar and the neighboring lateral geniculate nucleus (LGN). The local contrast and image salience models yielded highly correlated variance-explained maps (r = 0.8; Supplementary Figure 2). We found that these models best accounted for BOLD responses in the LGN (green outline) and inferior-lateral portions (light blue) of the pulvinar (Figure 2A, 2B; Supplementary Figure 3). This finding aligns with previous human neuroimaging studies demonstrating these regions’ sensitivity to basic visual patterns, such as flickering checkerboards (Arcaro et al., 2015; Cotton & Smith, 2007; DeSimone et al., 2015). In contrast, high-level feature processing was localized to medial and posterior regions of the pulvinar across subjects. The body and face pRF models best accounted for BOLD responses in these regions (Figure 2A&B; Supplementary Figure 3). The high correlation between face and body variance-explained maps (r = 0.93) may reflect the frequent co-occurrences of these features in natural scenes. The foreground pRF model showed a broader pattern of responsiveness, accounting for variance in both low-level and feature-responsive regions. Foreground maps were correlated with image contrast (r = 0.6) and salience (r = 0.65) as well as with face (r = 0.41) and body (r = 0.35) feature maps. As an important control comparison, not all features were linked to activations in the pulvinar.

**Figure 2.**
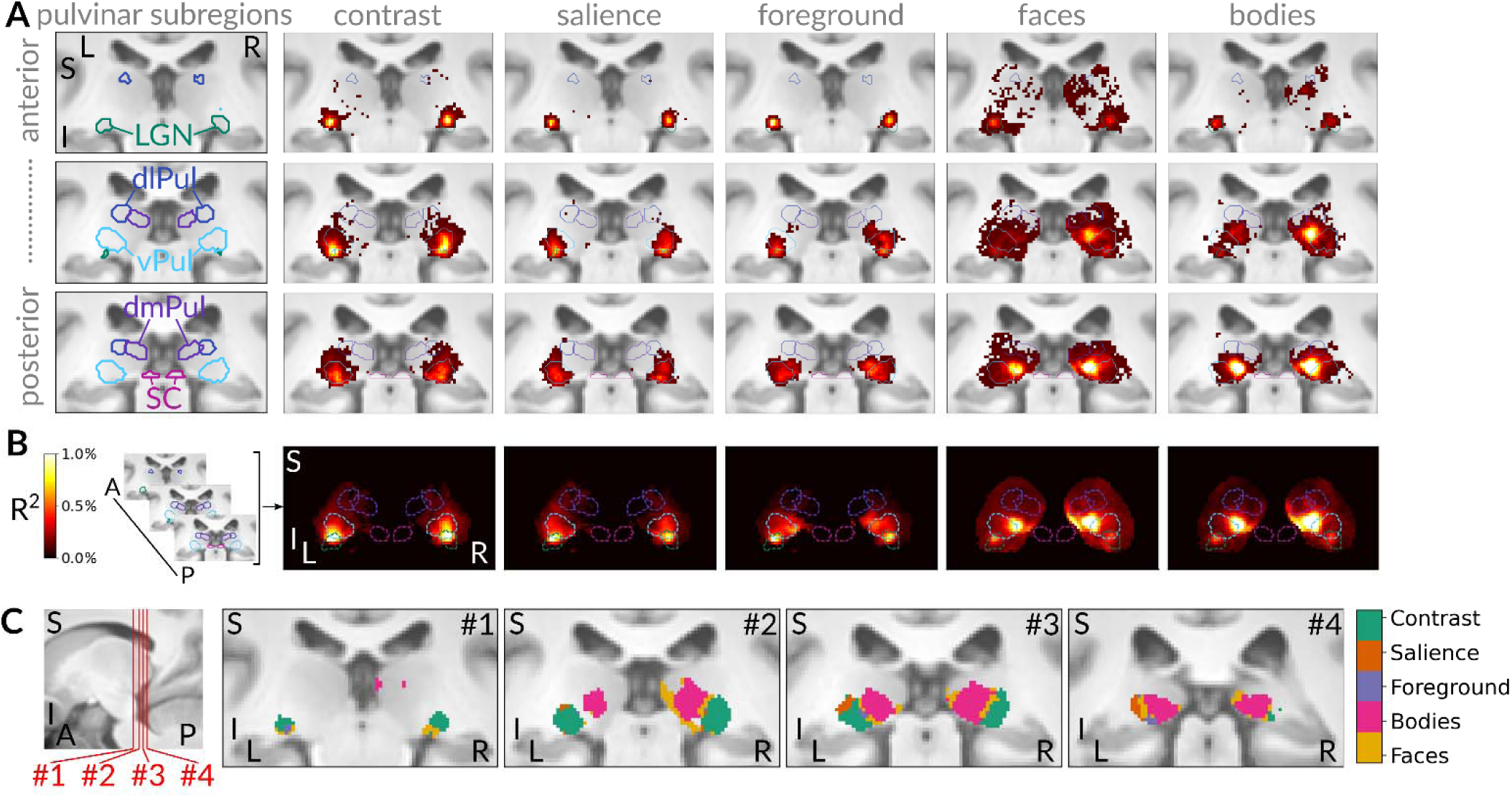
Distinct regions of the pulvinar responsive to low- and high-level visual features. A) Group-average (n = 8) variance-explained maps for six pRF feature models shown in three adjacent coronal images covering the LGN and pulvinar. The first column shows colored contours of five functionally defined subcortical ROIs derived from independent data overlaid on corresponding MNI anatomical coronal slices (green LGN; light blue ventral pulvinar dark blue dorsolateral pulvinar; dark purple dorsomedial pulvinar; magneta superior colliculus; see Methods for details; Arcaro et al. 2015; 2018). Remaining columns show variance-explained maps for each pRF model. Only voxels with > 0.2% variance explained are shown. B) Maximum intensity projection of variance-explained results along the anterior-posterior axis for each feature model. Dashed colored contours indicate the five ROIs. Variance-explained values from voxels outside the thalamus ROI were zero to 0%. C) Left, sagittal view of anatomy with red lines indicating which slices are plotted in subsequent panels. Right, coronal views of winner-take-all analysis, showing which feature model explained the most variance for each voxel within the posterior thalamus ROI. Only voxels with > 0.2% variance explained by any feature model are included.

Background and word pRF models (not shown), explained little variance in pulvinar activations (Supplementary Figure 4), and we therefore do not examine these models further.

A winner-take-all analysis comparing the variance explained by different feature models further illustrates this feature segregation within the pulvinar. Activity in the inferior-lateral pulvinar was best explained by image contrast and salience, while activity in the posterior-medial pulvinar was best explained by bodies and faces (Figure 2C). Notably, none of these models captured substantial variance in dorsal parts of the pulvinar associated with attentional filtering (Arcaro et al., 2018), suggesting that object-recognition processes are specifically linked to the ventral pulvinar. These results demonstrate a clear functional substructure within the pulvinar, with anatomically distinct regions exhibiting responses to low- and high-level features in visual stimuli. This organization parallels the hierarchical structure of feature selectivity observed in ventral visual cortex.

### Organized spatial coding of low- and high-level visual features within the pulvinar

Having identified distinct pulvinar regions responsive to low- and high-level visual content in real-world scenes, we next examined their spatial coding. Each pRF feature model produced a set of parameter estimates characterizing voxel-wise spatial selectivity throughout the pulvinar (Supplementary Figure 4). We focused on the image contrast and body pRF models, as these models explained the most variance in responses to the NSD stimuli within the lateral and medial pulvinar, respectively (Figure 2C).

With respect to low-level features, the contrast model revealed a bilateral topographic organization of contralateral visual space within inferior-lateral portions of the pulvinar across subjects (Figure 3B). The upper visual field (Figure 3B, second row; red) was represented ventral-laterally, while the lower visual field (blue) was represented superior and medially. Between these vertical meridian representations, there was a gradual progression of preferred visual angle crossing the horizontal midline (green). These representations were predominantly within central visual space (Figure 3B, third row) and exhibited strong lateralization to the contralateral visual field (Figure 3B, fourth row). At the group-average level, 73% of voxels having at least 0.1% variance explained had pRFs tuned within 3° eccentricity (range at individual subject level: 49–79%). pRF organization was consistent across individual subjects (Figure 3D) and aligns with prior electrophysiological work in non-human primates (Bender, 1981; Gattass et al., 1978) and fMRI studies in humans (Arcaro et al., 2015; DeSimone et al., 2015) showing an inverted retinotopic map in this region of the pulvinar. These results demonstrate that the spatial coding of the ventral pulvinar can be effectively probed by modeling responses to low-level feature content in real-world scenes.

**Figure 3.**
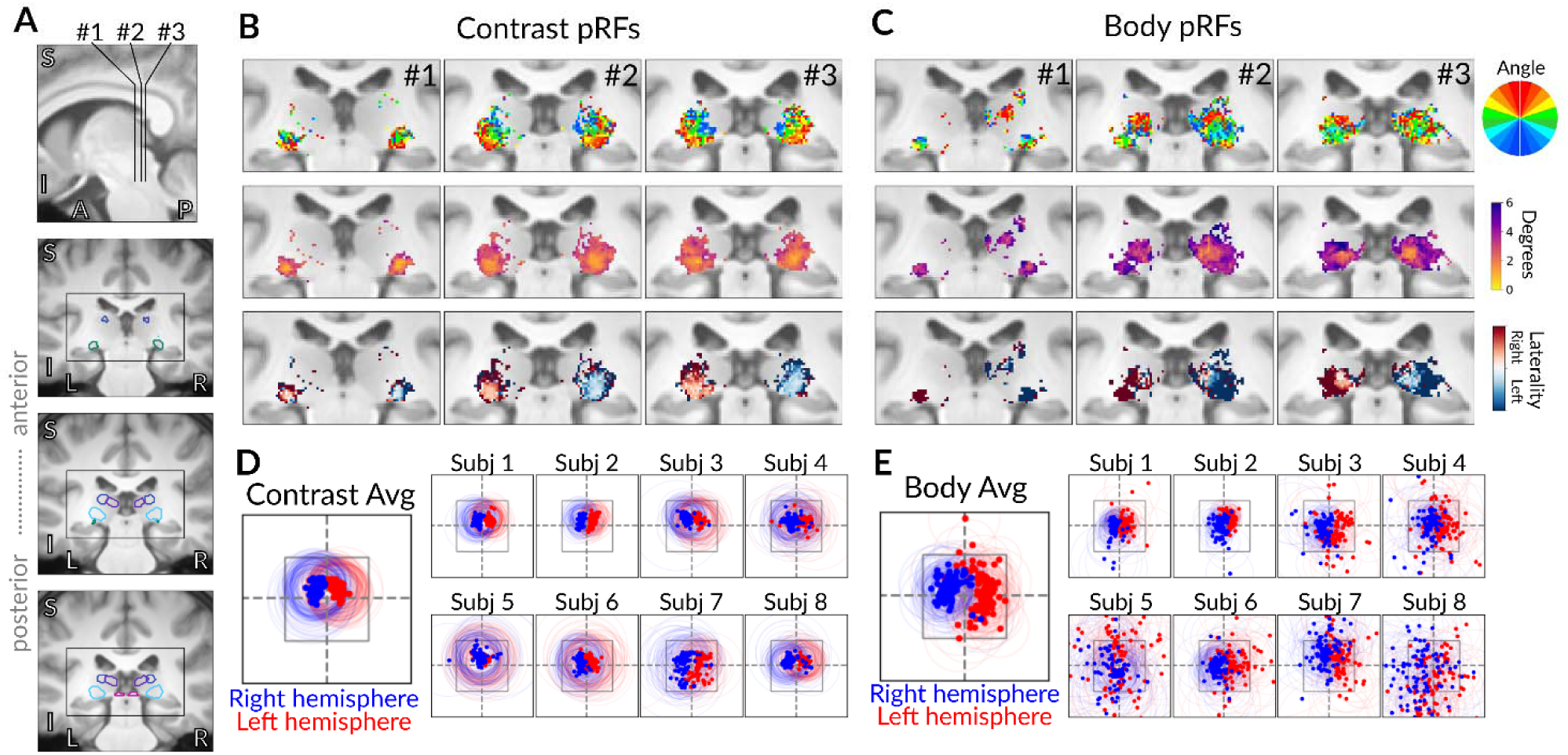
Spatial coding of low- and high-level visual features in the pulvinar. A) An MNI anatomical reference illustrating the location and extent of the coronal slices used in panels B and C. Colored contours indicate the five subcortical ROIs shown in Figure 2. B) Group-average (n = 8) results from the contrast pRF model. From top to bottom, polar angle, eccentricity, and visual-field laterality are shown for three coronal sections. Data were thresholded at 0.1% variance explained (by the contrast pRF model). C) Group-average results from the body pRF model. Same format as panel B (with data thresholded based on body pRF model results). D) Visual field coverage plots for the contrast pRF model. pRFs from the top 100 voxels in each hemisphere (in terms of variance explained) are plotted on the visual field. Dots indicate the centers of pRFs while circles indicate the extent of the pRFs (+/-2 standard deviations). The gray square indicates the size and location of the stimulus. Dashed lines indicate the vertical and horizontal meridians. The larger subpanel shows results from a group average (i.e., based on averaging pRF parameters across participants in each voxel), while the other subpanels show results from individual subjects. E) Visual field coverage for the body pRF model, in the same format as panel D.

Surprisingly, responses to high-level visual content also exhibited clear topographic organization. The body pRF model revealed bilateral representations of contralateral visual space in pulvinar regions posterior and medial to the contrast model activations (Figure 3C, second row). Upper visual field representations were localized to dorsal medial portions, while horizontal meridian and lower visual field representations were ventral and lateral. The medial pulvinar contained a central visual field representation surrounded by peripheral representations (Figure 3C, third row). Body pRFs were generally lateralized to the contralateral visual hemifield (Figure 3C, fourth row) and covered the extent of the visual field stimulated by the NSD images (Figure 3E). At the group-average level, only 19% of voxels having at least 0.1% variance explained had pRFs tuned within 3° eccentricity (range at individual level: 17–57%). These results demonstrate that the processing of high-level categorical content in the pulvinar is anchored to spatial coding of sensory input.

Comparison of visual field representations derived from the contrast and body models revealed two distinct, mirror symmetric retinotopic zones. The organization of visual space along the polar angle dimension for the body model was inverted relative to that in the lateral pulvinar for the contrast model. The lower visual field representations of both models overlapped (Figure 3F; Supplementary Figure 5, dashed lines), whereas the upper visual field representation from the contrast model was located ventrolaterally and that of the body model dorsomedially. This offset and inversion of the visual field between models indicates that lateral and medial regions of the pulvinar contain distinct maps of visual space.

### Model-free analyses reveal similarity in representations between cortex and pulvinar

As a complement to our model-based analyses, we devised a model-free approach to relate cortical and subcortical representations of natural scenes. Prior research has shown that different regions of the pulvinar are anatomically connected (Shipp, 2003) and functionally coupled (Arcaro et al., 2018) with different parts of visual cortex. The anatomical locations of the contrast- and body-selective subregions align with evidence for a gradient of pulvino-cortical connectivity (Arcaro et al., 2015; Shipp, 2003), suggesting that visually evoked activity in different parts of the pulvinar might resemble activity in functionally corresponding cortical regions. To test this hypothesis and the functional specificity of such connectivity, we conducted a correlational analysis between the pulvinar and cortex based on activations from the NSD dataset.

We analyzed single-trial BOLD response amplitudes in the pulvinar and cortex, derived using GLMsingle to capture the response of each voxel to individual image trials in NSD (Figure 4A; see Methods for details). For each subject, we identified pulvinar voxels showing the highest variance explained by the contrast and body models. We then computed trial-by-trial correlations between the BOLD responses in these pulvinar voxels and all cortical voxels (Figure 4B). This analysis revealed robust correlations between the pulvinar and extensive portions of the visual cortex for both the contrast-peak and body-peak pulvinar voxels, with weaker correlations in non-visual cortex (Figure 4C). Contrast-peak voxels showed strong correlations with both early and anterior visual cortex, whereas body-peak voxels correlated more selectively with anterior visual cortex. These findings demonstrate distinct patterns of cortical coupling for lateral and medial regions of the ventral pulvinar.

**Figure 4.**
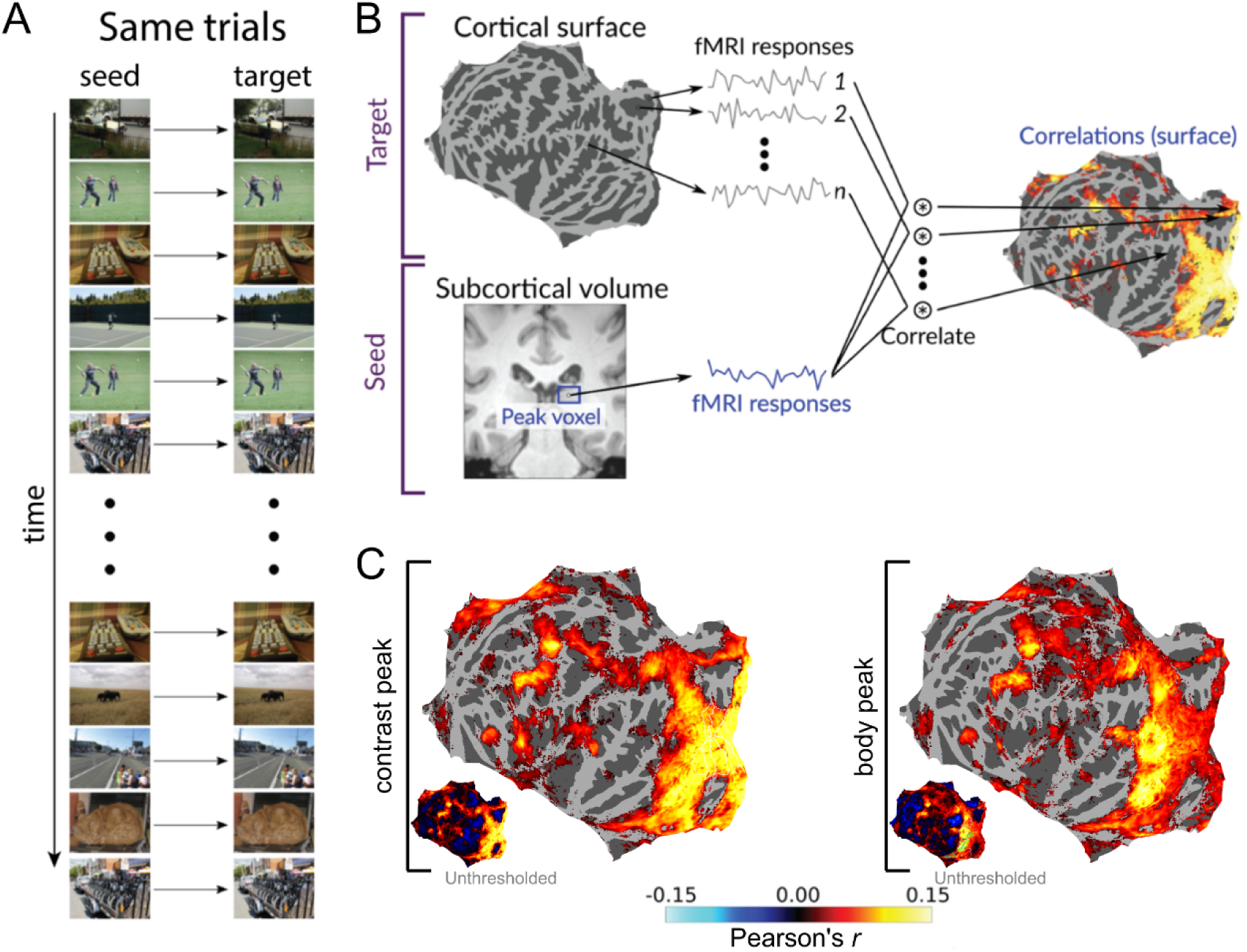
Contrast- and body-selective pulvinar subregions are co-active with distinct cortical regions. A) Schematic showing the same-trial correlation analysis, where correlation is computed between BOLD responses observed at the same time. B) Schematic showing how subcortical-to-cortical correlations are computed. For a given seed voxel in the pulvinar, we correlated the activity in that seed voxel with activity in each vertex on the cortical surface, resulting in a map of correlations across the cortical surface. C) Cortical correlations resulting from seeding the peak voxel in the pulvinar for the (left) contrast and (right) body models. Data are thresholded based on a bootstrap procedure (see Methods for details). Inset images show unthresholded maps.

To isolate correlations driven by stimulus-evoked responses from other potential sources of co-activation (such as arousal and attention), we recomputed correlations between the pulvinar and cortex using responses to the same image but from different trials (Figure 5A). This approach yielded more spatially localized correlations. For the contrast peak, cortical correlations were predominantly confined to posterior occipital cortex, aligning with the extent of early visual retinotopic maps (V1-hV4) identified from a separate localizer experiment (Figure 5B; see Methods for details). Body peak co-activations were primarily constrained to extrastriate visual cortical areas responsive to faces and bodies, including EBA, FBA, and FFA (Figure 6C). Notably, the body peak showed no correlation with early visual cortex or other category-selective extrastriate areas, such as place (e.g., TOS, PPA) and word areas (e.g., OWFA, VWFA). These findings demonstrate functionally specific coupling between pulvinar and cortex in response to real-world stimulation, reflecting an alignment of processing between the thalamus and cortex.

**Figure 5.**
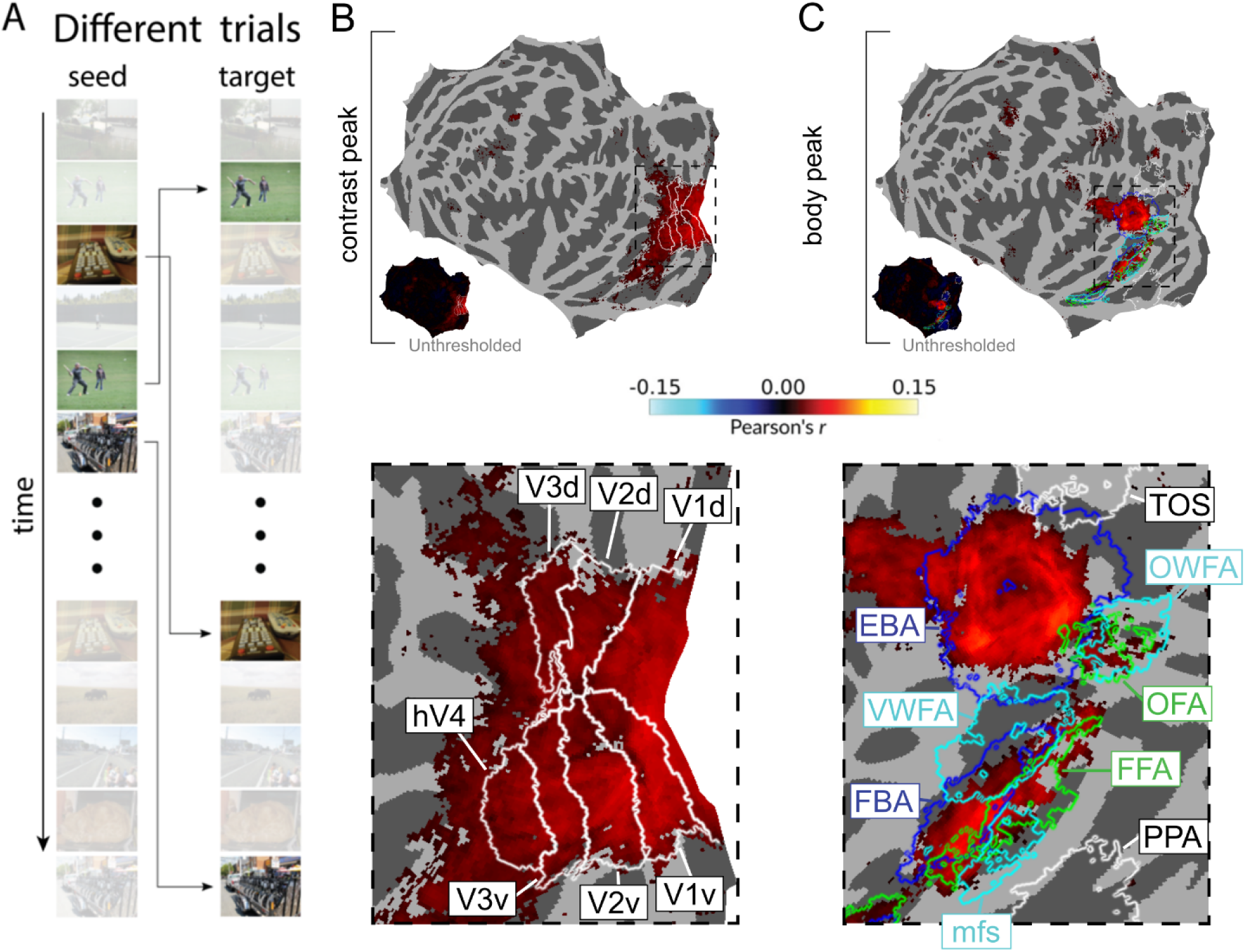
Different-trials analysis reveals highly specific pulvino-cortical correspondences. A) Correlation is computed between BOLD responses evoked by the same stimulus image but on different trials. B) Cortical correlations corresponding to the peak pulvinar voxel for the contrast pRF model. Data are thresholded based on a bootstrap procedure (see Methods for details). Bottom image shows zoomed-in view of peak correlations within occipital cortex and correspondence to cortical ROIs defined based on separate localizer experiments (see Methods). C) Cortical correlations corresponding to the peak pulvinar voxel for the body pRF model. Same conventions as B.

**Figure 6.**
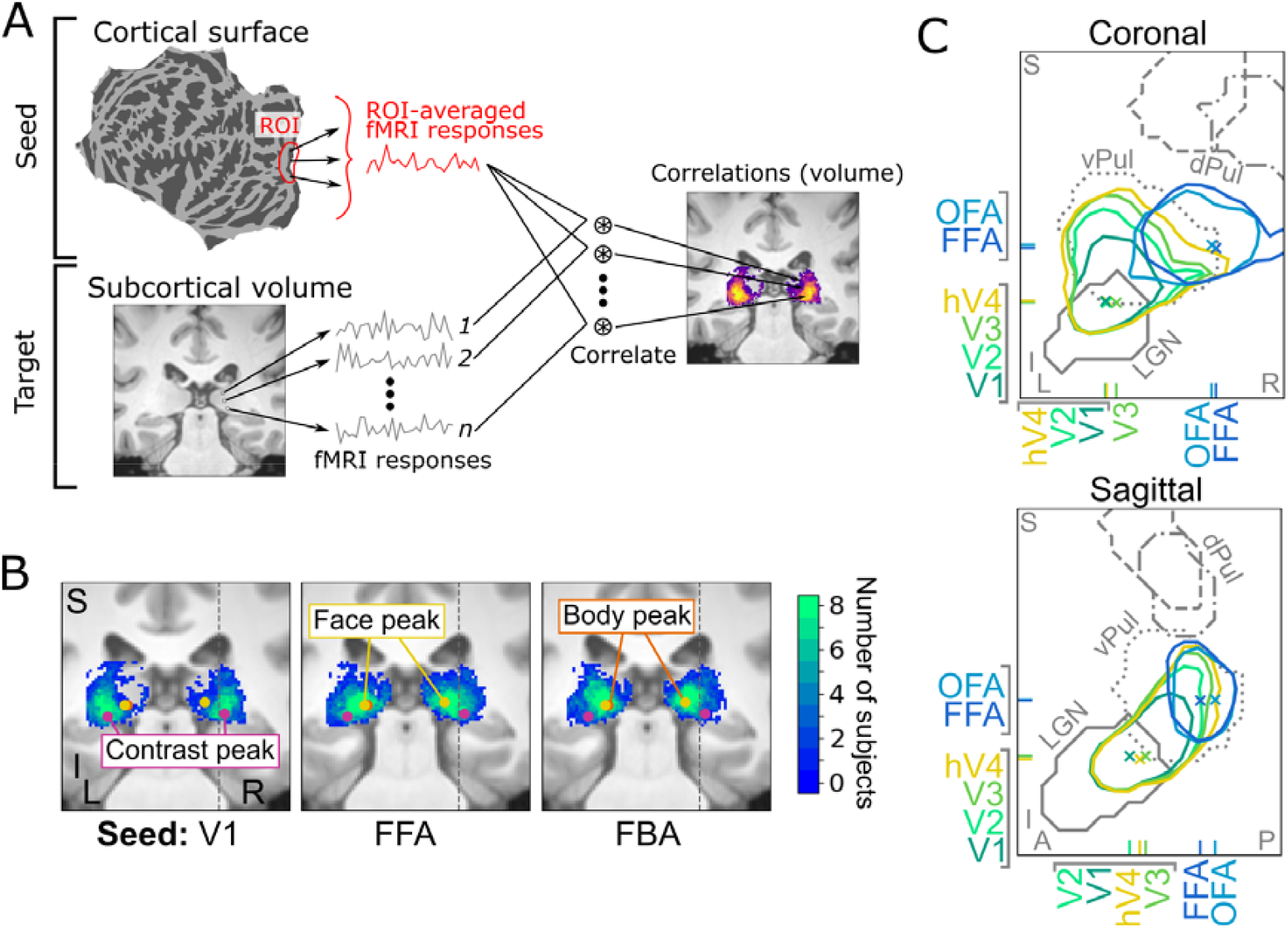
Progression along ventral cortical hierarchy recapitulated in the pulvinar. A) Schematic showing how cortical-to-subcortical correlations were computed. For a given cortical surface ROI, we correlated the ROI-averaged cortical activity with activity in each subcortical voxel, resulting in a volumetric correlation map throughout the pulvinar. Correlations were computed on BOLD responses to the same stimulus across different trials. B) Coronal slices showing pulvinar correlation maps corresponding to V1, FFA, and FBA cortical ROIs. At each subcortical voxel, we plot the number of subjects for whom correlation values passed a threshold of |r| > 0.01 (see Methods for details). We also label the peak model-based variance-explained value for the contrast, face, and body models for the depicted slice. Faint gray dashed lines in the coronal slice indicate the same location across maps. C) Comparison of correlations across the pulvinar for visual areas V1, V2, V3, hV4, OFA, and FFA. Colored lines indicate half-max contours of the maximum intensity projection of cortical-to-subcortical correlation results for six cortical ROIs. Tick marks indicate the position of the maximum cortical-to-subcortical correlation value for a given cortical ROI. Gray lines indicate contours of maximum intensity projections of the anatomical ROIs from Figure 2B.

To contextualize these results with respect to the entire pulvinar, we performed a third analysis correlating responses from various cortical areas with responses from each voxel within the pulvinar (Figure 6A). As in Figure 5, this analysis was performed using responses to the same images from different trials. This analysis revealed that V1 correlations were strongest within the LGN and ventrolateral areas of the pulvinar (Figure 6B), consistent with the greater anatomical connectivity of those subcortical regions with early visual cortex (Shipp, 2003). In contrast, FFA and FBA correlations were biased toward superior and medial aspects of pulvinar (Figure 6B). Notably, voxels best explained by the contrast pRF model (Figure 6B; purple circles) aligned with the peak V1 correlation, while those best explained by the face (yellow circles) or body (orange circles) pRF models aligned with the peak FFA and FBA correlations.

Visualizing cortical correlation maps using half-max contours revealed a clear anterior-inferior-lateral to posterior-superior-medial progression as the cortical seed area moved along ventral visual cortex from V1 posteriorly to hV4 anteriorly (Figure 6C; see Methods for details). While all early visual cortical areas showed correlations with the LGN and ventrolateral pulvinar, the maximal superior-medial extent of correlation increased for each successive visual cortical area. This pattern suggests the existence of a hierarchical functional gradient throughout the pulvinar, mirroring the organization of the ventral visual cortex.

## Discussion

Our analysis of BOLD responses to tens of thousands of natural scenes revealed a complex functional architecture within the pulvinar. We identified distinct regions responsive to low-level visual features and high-level object categories, particularly faces and bodies. This functional clustering was embedded within a broader functional organization of the pulvinar that mirrors the topography of visual response properties observed across the ventral visual cortical hierarchy. Our findings suggest that pulvinar subregions may interact with corresponding levels of cortical processing, and that the pulvinar may play a more integral role in visual processing than previously appreciated.

### High-level visual processing in the pulvinar

While the pulvinar has long been associated with visual processing, most studies have focused on its involvement in low-level visual feature processing (Bender, 1981; Petersen et al., 1985) or in attention (Kastner et al., 2004) and motor processes (Wilke et al., 2010). Our results provide strong evidence for its involvement in high-level visual perception. We identified a focal region in the posterior medial pulvinar that selectively responds to images of faces and bodies, extending previously reported weak biases for such stimuli (Arcaro et al., 2018; Wen et al., 2023). This finding aligns with observations of responsiveness to face-like images (Nguyen et al., 2012) and facial emotion (Maior et al., 2010) in the macaque pulvinar.

A main contribution of the present work is a characterization of these high-level responses, demonstrating their specificity in relation to spatial coding, stimulus feature selectivity, localization within the pulvinar, and functional coupling with corresponding visual category clusters in cortex. Our results suggest that the pulvinar may play a more integral role in object recognition than previously thought, potentially contributing to the rapid and efficient processing of socially relevant stimuli like faces and bodies (Pessoa & Adolphs, 2010). Converging with recent work showing face responses in the superior colliculus (Yu et al., 2024), our results indicate a broader role for subcortex in high-level vision.

Distinct from this category selectivity, we confirmed previous findings of low-level visual feature processing in the ventral lateral pulvinar (Bender, 1981; Petersen et al., 1985), reflecting its anatomical connections with early visual areas (Shipp, 2003). These results indicate that the pulvinar is involved in multiple states of object processing, from low-level feature extraction to high-level category representation.

### A thalamo-cortical hierarchy of object processing

The face and body clusters identified were embedded within a broader functional organization of the pulvinar. Between the regions responsive to low-level visual features (e.g., local contrast) and high-level visual features (e.g., category labels), we found preferential responsiveness to foreground images and image saliency, indicating sensitivity to intermediate-level visual information. This functional differentiation within the pulvinar parallels the hierarchical organization of the ventral visual cortex.

Our results suggest that the pulvinar is not a separate, independent subcortical pathway for object processing. Indeed, the hierarchical organization of thalamo-cortical connections suggests that the pulvinar could be viewed as an integral part of the ventral visual cortical pathway. The pulvinar has limited internal connectivity (Imura & Rockland, 2006) with most of its connections being with cortex. Prior studies have revealed a gradient of anatomical (Adams et al., 2000; Baleydier & Morel, 1992; Webster et al., 1993) and functional (Arcaro et al., 2018) connectivity across the pulvinar, moving from posterior to anterior visual cortex. Converging with these prior studies, our analysis of processing during natural scene viewing revealed a gradient of functional coactivation between the pulvinar and cortex that mirrors the hierarchical organization of the ventral visual pathway.

What functional role could this pattern of connectivity with cortex serve? We speculate that the hierarchical organization of connectivity may reflect the pulvinar’s broader role in gating cortical visual processing and synchronizing cortico-cortical communication via cortico-thalamic loops. Studies in macaques have shown that pulvinar neurons can modulate the gain of visual responses in cortex (Purushothaman et al., 2012) and facilitate communication between distant cortical areas (Saalmann et al., 2012). Whereas most work pursuing this hypothesis has focused on understanding the pulvinar’s role in cortical attention networks (Fiebelkorn et al., 2019; Saalmann et al., 2012; Zhou et al., 2016), our results provide some of the first evidence that the pulvinar may also play an important role in regulating cortical brain networks dedicated to object vision. These influences occur at multiple stages of the cortical hierarchy (De Souza et al., 2020; Sherman, 2017), potentially integrating diverse visual information and facilitating the binding of relatively basic visual features into coherent object representations.

This hierarchical integration of the pulvinar contrasts with recent findings of rapid face responses in the superior colliculus (Yu et al., 2024). While both subcortical structures contribute to high-level vision, they exhibit complementary mechanisms. The pulvinar interfaces with multiple stages of the ventral visual hierarchy, potentially integrating information across different levels of representation and facilitating more complex object and category representations. In contrast, the superior colliculus generates a rapid signal, potentially derived from V1 input, that may facilitate quick orienting towards faces. These distinct, yet complementary roles highlight the diverse subcortical contributions to object recognition and perceptual processing.

The functional hierarchy we observed within the pulvinar also supports the existence of multiple thalamo-cortical pathways, consistent with previous work showing distinct functional connectivity patterns between pulvinar subregions and ventral and dorsal cortices (Arcaro et al. 2018). Similarly, studies in macaques have highlighted the pulvinar’s contributions to both dorsal and ventral visual processing streams (Kaas & Lyon, 2007; Shipp, 2003). The complex functional architecture revealed by our study underscores the pulvinar’s potential as a key node in distributed networks supporting varied visual functions. While our results emphasize the integration of processing within the ventral visual pathway, it remains an open question the extent to which the pulvinar might integrate information across multiple processing streams, and even potentially serve as a hub for multi-sensory integration (Froesel et al., 2021).

### Retinotopy as an organizing principle of the visual pulvinar

A striking feature of our results is the prominence of retinotopic organization at multiple levels of visual responsiveness in the pulvinar, including regions selective for high-level visual object categories. This preservation of retinotopic organization mirrors findings in inferotemporal cortex, where spatial sensitivity persists (Op De Beeck & Vogels, 2000) even in face- and object-selective areas (Arcaro et al., 2020; Hasson et al., 2002; Kay et al., 2015). This consistent spatial coding across cortical and subcortical structures suggests a fundamental role for retinotopy in visual processing that extends beyond early visual processing areas, providing a framework for integrating diverse information within a coherent spatial reference frame, from basic feature detection to complex object representations.

These results align with proposals that retinotopic organization plays a crucial role in the development and organization of visual cortex, potentially serving as a scaffold for the emergence of more complex visual representations (Arcaro & Livingstone, 2021). Interestingly, pulvinar connectivity with cortex emerges early in gestational development (Shatz & Rakic, 1981), exhibiting a hierarchical and retinotopic organization at birth (Ayzenberg et al., 2025), which precedes the formation of functional clustering in cortex such as face-(Livingstone et al., 2017) and word-selectivity (Dehaene-Lambertz et al., 2018). This developmental timeline suggests that the pulvinar may play a critical role in shaping the ventral visual cortical hierarchy (Bourne & Rosa, 2006), potentially guiding learning and functional specialization in cortex (O’Reilly et al., 2021).

### Towards studying visual networks with higher ecological validity

Our study demonstrates the utility of using naturalistic stimuli to probe processing throughout the visual system. While naturalistic stimuli are often regarded as too uncontrolled for systematic investigation, we have shown that computational approaches can be used to quantify the contribution of various aspects of visual processing, from low-to high-level visual properties, and link this processing with specific neural circuits. Our findings isolating regions of the ventral lateral pulvinar selective to local image contrast content in the real-world scenes converges with prior work using less naturalistic stimuli, serving as a validation of this approach and demonstrating a principled way to explore neural responses to basic visual properties within real-world contexts.

The use of naturalistic stimuli in conjunction with computational modeling approaches offers several advantages. First, it allows for the investigation of visual processing under conditions that more closely approximate real-world vision. Second, it enables the simultaneous examination of multiple levels of visual processing, from low-level features to high-level object categories. Third, it may reveal functional properties that are not easily observable with more constrained stimuli, as evidenced by our discovery of robust visual-category selectivity in the pulvinar. The success of this approach in uncovering novel aspects of pulvinar function underscores its potential for embracing complexity of natural visual input. This shift towards more ecologically valid paradigms promises to deepen our understanding of how the brain processes the rich visual world.

## Conclusion

Our study demonstrates that the pulvinar contains multiple retinotopic representations of the visual scenes, ranging from low-level features to high-level object categories. These representations are localized to distinct anatomical regions that align well with known patterns of pulvino-cortical anatomical connectivity. Our results support the hypothesis that the pulvinar plays an integral role in coordinating communication between visual cortical areas across multiple levels of processing.

These findings underscore the importance of considering subcortical contributions to object vision. Unravelling the complexities of cortical visual processing, such as in face and body recognition, may require a more comprehensive understanding of their interactions with subcortical structures like the pulvinar and superior colliculus (Yu et al., 2024). Our work provides a foundation for future investigations into the computational roles of the pulvinar in object recognition and other high-level visual processes.

## Acknowledgements

This work was supported by NSF CRCNS IIS-1822683 (K.N.K.) and by NSF NRT-UtB1734815 (D.R.G., as trainee).

## Data and code availability

Data used in this paper are available online at http://naturalscenesdataset.org/. Code used to perform the analyses and generate the figures are available online at https://github.com/guestdaniel/Guestetal2025_NSDPulvinar.

## Author contributions

D.R.G. performed the analyses, produced the figures, and wrote the original draft. E.J.A. collected the original neuroimaging data (see Allen et al., 2022 for details), edited the paper, and assisted with figure design. K.N.K. designed and supervised the original neuroimaging experiments and analyses (see Allen et al., 2022 for details), edited the paper, and co-supervised the project (analysis, figures, and writing). M.J.A. provided ROI labels, edited the paper, and co-supervised the project (analysis, figures, and writing).

## Methods

### Participants

Eight participants (six females and two males; age range, 19-32 years) participated in the NSD study, which was approved by the Institutional Review Panel of the University of Minnesota. All participants provided informed consent and had normal or corrected-to-normal acuity. For detailed participant information, see Allen et al. (2022).

### Dataset

The NSD comprises fMRI measurements from 8 participants viewing 9,000-10,000 distinct natural color scenes (22,000-30,000 trials) over 30-40 scan sessions. Scanning was conducted using a 7T MRI scanner, with whole-brain gradient-echo EPI at 1.8-mm isotropic resolution and 1.6-s repetition time. Images were sourced from the Microsoft Common Objects in Context (COCO) database (Lin et al., 2015), square cropped, and presented at 8.4° x 8.4° visual angle. Stimulus images reproduced here (Figures 1, 4, 5 and S1) are thus modifications of the original images used in compliance with COCO database’s Creative Commons 4.0 license (https://creativecommons.org/licenses/by/4.0/). A set of 1,000 images was shared across participants, with the remaining images unique to each participant. Images were presented for 3 s each, with 1-s gaps between them.

Data preprocessing included temporal interpolation for slice time correction and spatial interpolation for head motion correction and compensation for spatial distortion. Single-trial beta weights, representing BOLD response amplitudes, were estimated using a general linear model. The study utilized “Version 3” trial response estimates (Allen et al., 2022), which incorporate voxel-specific hemodynamic response functions, denoising using GLMdenoise (Kay, Rokem, et al., 2013), and ridge regression for trial responses estimation. For cortical surface data, we utilized the NSD-prepared subject-native surface data, registered and transferred to the fsaverage template using nearest neighbor interpolation. For subcortical data, we used the 1.0-mm volumetric preparation of the NSD data in subject-native space, mapped into the MNI template space using the provided T1-to-MNI transformation. To control for inter-session variability, all data were z-scored within each voxel (or vertex for surface preparations) on a session-by-session basis.

### Regions of Interest (ROIs)

Subcortical analyses focused on a manually defined region of the posterior thalamus that included the pulvinar, lateral geniculate nucleus (LGN) and superior colliculus (SC). This region was identified on each participant’s native-space T1-weighted anatomical image, using the contrast between gray and white matter (for more details, see Arcaro et al. 2015). To ensure consistency across subjects, each subject’s subcortical ROI was then transformed to 1mm MNI space using nearest-neighbor interpolation. A group average was constructed by selecting all voxels labeled as posterior thalamus in at least 3 out of 8 participants, then projected back to each participant’s native 0.8-mm anatomical space.

Specific structures within this ROI were identified based on previously reported group average functional data (Arcaro et al., 2015). These data were warped from 1-mm MNI space to each participant’s native space. The ventral pulvinar was defined by voxels in “vPul1” or “vPul2” retinotopic maps. Dorsomedial and dorsolateral pulvinar were defined by correlations with the precuneus and retinotopic portions of the frontoparietal cortex (IPS1-5, SPL1, FEF, and IFS), respectively. The LGN and SC were defined by their respective retinotopic maps. ROIs were manually edited to exclude voxels outside the anatomical extent of each structure and transformed back to 1mm MNI space for group-average ROI construction.

### Stimulus feature maps

Feature maps for image contrast, saliency, foreground, background, and specific image categories (faces, bodies, and words) were generated for fMRI analyses. Local image contrast maps were computed by converting images to greyscale, resizing to 800 × 800 pixels, squaring to approximate luminance response, and calculating local contrast within a 51×51 grid (grid element size: 0.168 × 0.168 degrees). The grid was padded with half a grid element on all sides to capture transitions between the image and the mean grey background (for more detail, see Allen et al., 2022). Saliency maps were generated using a pre-trained deep neural network model (Kroner et al., 2020) trained on the SALICON dataset, a fixation dataset based on the COCO image database (Jiang et al., 2015). As the NSD images also were sourced from the COCO database, this model was expected to generalize well to the NSD stimuli. Saliency maps were cropped and resampled to match the NSD stimuli from the COCO database (Allen et al., 2022). All other feature maps comprised binary mask annotations. Face annotations were generated using the RetinaFace model (https://github.com/deepinsight/insightface/tree/master/RetinaFace), which identifies rectangular bounding boxes around faces. Word annotations were created using the EAST text detector (https://github.com/argman/EAST), which outputs bounding quadrangles or rectangles around text. Quadrangles were converted to enclosing rectangles for consistency with other annotations. Whole body annotations were sourced from the Microsoft COCO dataset (http://images.cocodataset.org/annotations/annotations_trainval2017.zip), using bounding boxes for human bodies (category ID 1) and animals (categories 16:25). Foreground annotations were derived from segmentations across all 80 object categories, and background annotations were defined as the inverse of the foreground annotations.

### Population receptive field (pRF) analysis

A pRF encoding model with compressive spatial summation was employed to characterize spatially specific responses to stimulus feature maps within each subject’s subcortical ROI. The model was implemented using a modified version of AnalyzePRF (http://cvnlab.net/analyzePRF/), which estimates voxel-wise parameters for each feature map (Kay, Winawer, et al., 2013; see Allen et al., [2022] for details), including spatial position, pRF size, gain, and variance explained. Separate models were fit for each feature map, with variance explained (R^2^) reflecting the percentage of variance in trial-averaged BOLD responses accounted for by the model. Only stimuli unique to each subject were included, excluding the common stimuli viewed by all participants. For non-contrast and non-saliency pRF models, a baseline term was added to account for responses to image regions covered by the annotated feature. For group-level analysis, voxel-wise parameter estimates were transformed to 1mm MNI space using linear interpolation, and median values then were calculated across participants for all parameters.

### Winner-take-all pRF analysis

The performance of different features was evaluated with a winner-take-all analysis. Variance explained values for each feature model were averaged across participants in the shared 1mm MNI space (see above). Then, each voxel within the thalamus ROI with at least 0.2% variance explained by any feature was labeled to indicate which feature model explained the most variance.

### Pulvino-cortical correlation analysis

Correlation analyses comprised two parts: (1) correlating single-voxel pulvinar responses with those across the entire cortical surface, and (2) correlating average responses for cortical ROIs with all pulvinar voxels.

For the first analysis, we identified voxels with the highest variance explained for the contrast model (“contrast peaks”) and for the body model (“body peaks”) in each participant’s native space. Trial response estimates from these voxels were correlated with response estimates from every vertex on the fsaverage cortical surface. This process was repeated 1000 times for each participant with trials randomly resampled with replacement (i.e., bootstrapped). For visualization of results on the cortical surface (Figs 4, 5), the correlation coefficient, averaged across bootstrap replicates and participants, was shown only if at least 95% of the bootstrapped correlation coefficients exceeded a value of 0 in a majority of participants in the given vertex.

For the second analysis, we constructed a set of cortical ROIs covering most of the visual cortex. These included: (1) retinotopic areas (V1, V2, V3, and hV4) defined from the pRF localizer experiment included with NSD (Allen et al., 2022), (2) additional retinotopic areas from the Wang probabilistic retinotopic atlas (Wang et al. 2015), and (3) category-selective areas (OFA, FFA, aTL-faces, EBA, FBA, PPA, and VWFA) from the NSD functional category localizer experiment (Allen et al., 2022). ROIs were pruned on an individual-participant basis to avoid overlap. Average responses were generated for each cortical ROI in fsaverage space and correlated with the trial response estimates from voxels within the subcortical ROI.

We performed analyses between the pulvinar and cortex using both same-trial and different-trial (for same image) pairings of response estimates. For same-trial correlations, each response in the subcortex was correlated with responses from the same trials in the cortex. For different-trial correlations, we leveraged that each participant viewed each distinct image three times. This allowed us to identify six possible pairs of image repeats, resulting in six response pairings for 10,000 images across different trials of the same image. We computed the correlation for each pairing, and averaged the resulting six correlation coefficients to derive a final correlation estimate between the pulvinar and cortex.

## Supplementary Figures

**Supplementary Figure 1.**
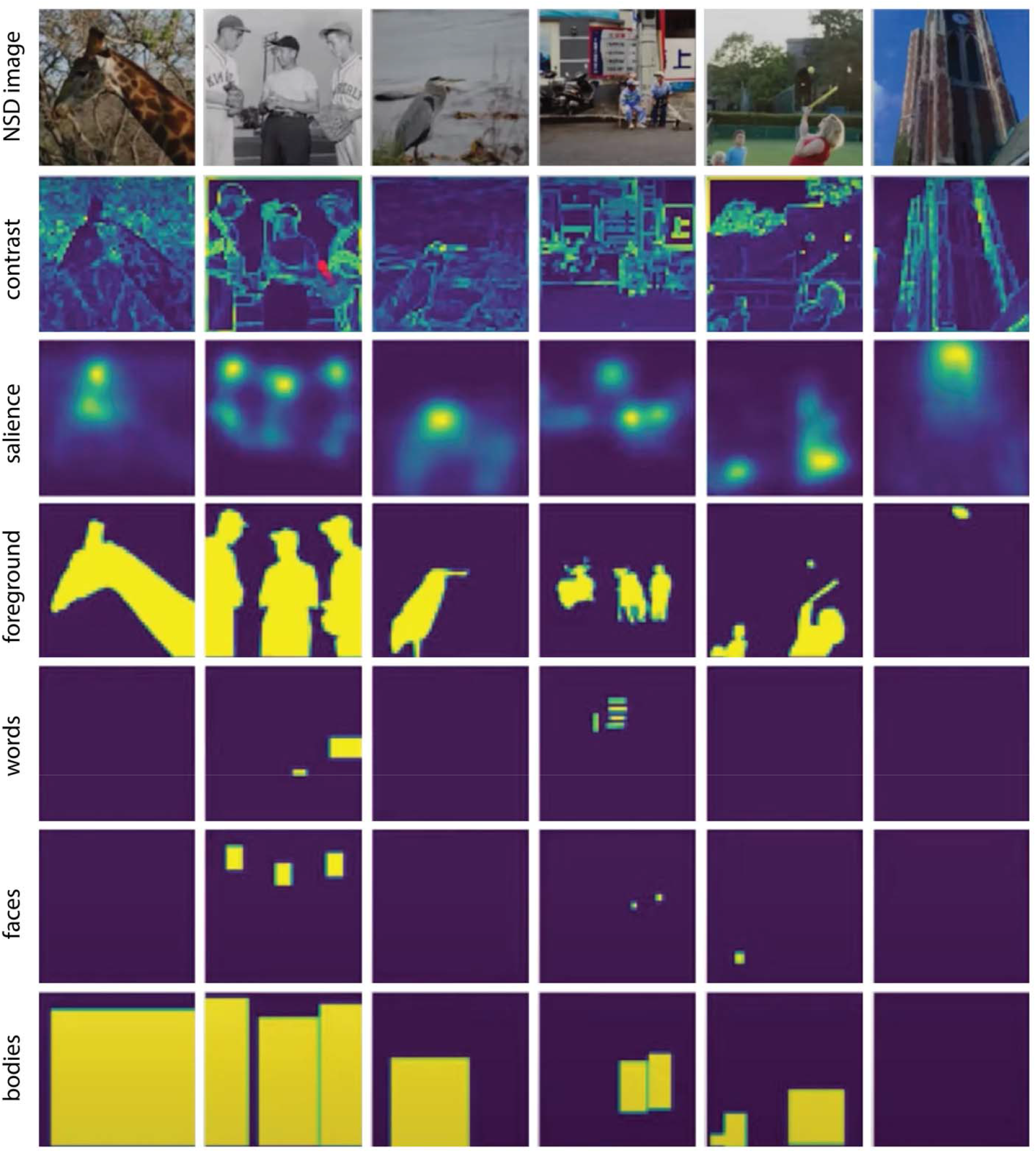
Examples of stimulus feature maps. Six example natural scene stimuli (columns) used in the fMRI experiment are shown along with the stimulus feature maps derived from these images for contrast, salience, foreground, word, face and body pRF encoding models (rows).

**Supplementary Figure 2.**
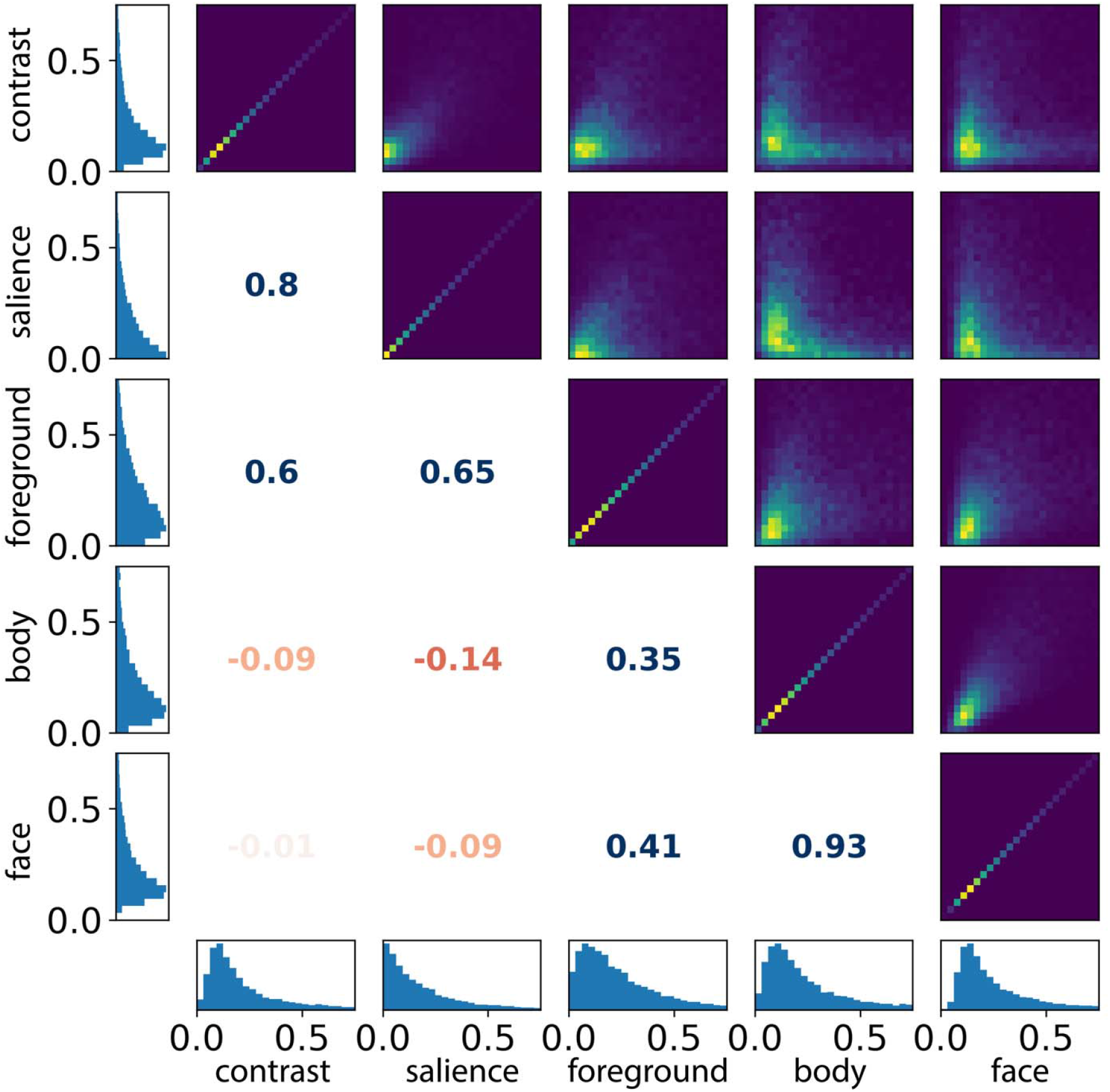
Histograms and correlation coefficients for voxel-wise variance-explained values (in percent) for pairs of different pRF models. Square panels on and above the main diagonal show 2D histograms, where brighter colors indicate more voxels with a given combination of variance-explained values for the two pRF models. Numbers indicate Pearson’s correlation coefficient between variance-explained values for the two pRF models (fonts are colored correspondingly, with red indicating negative correlations and blue indicating positive correlations). Rectangular panels on the edges indicate 1D histograms of variance-explained values for single pRF models.

**Supplementary Figure 3.**
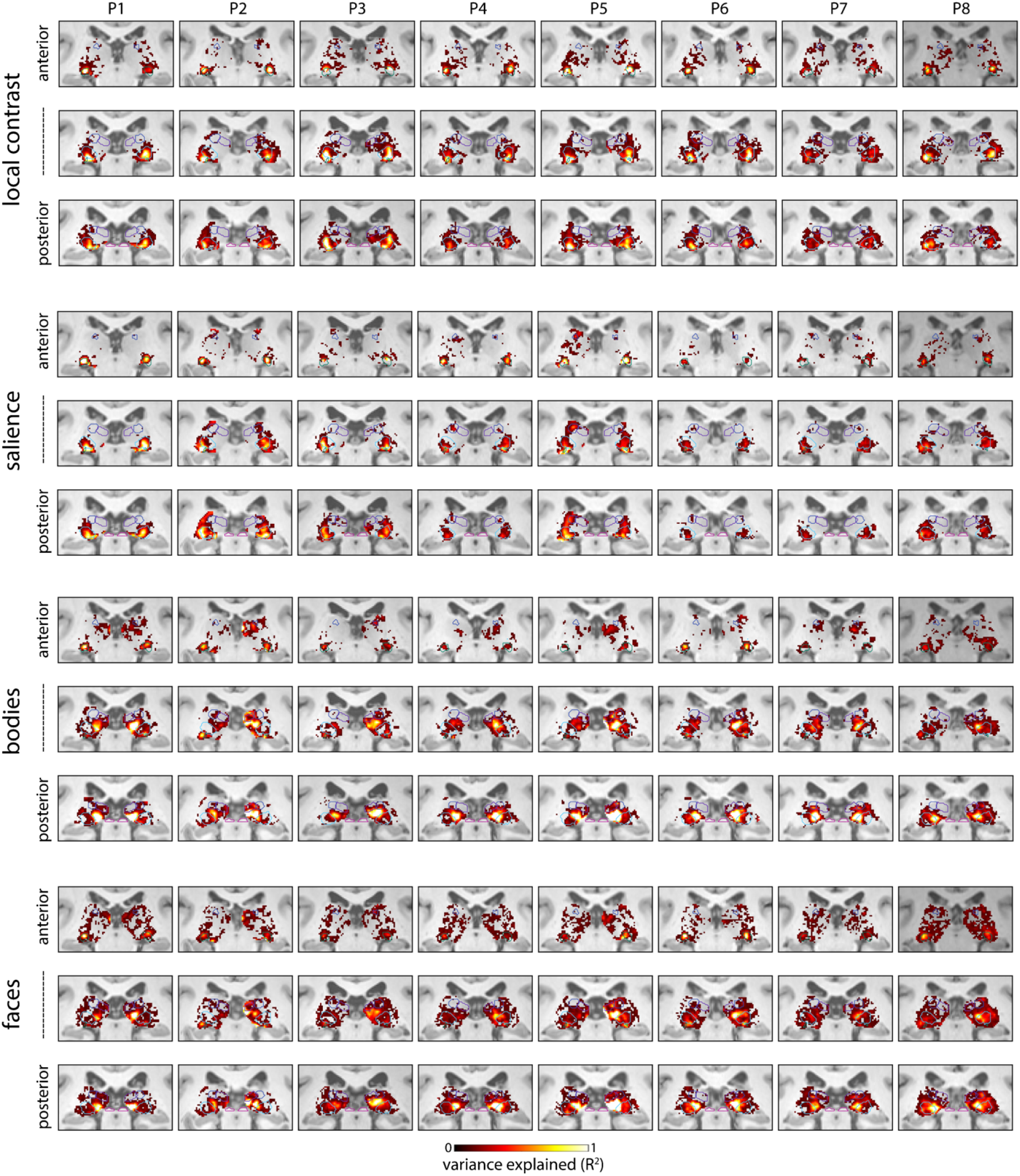
Individual-subject variance explained maps for the contrast, salience, bodies, and faces pRF models. The coronal slices shown are the same as in Fig. 2A; thin colored outlines indicate the five thalamus ROIs as in Fig. 2A. Only voxels with >0.2% variance explained are shown. See Figure 2 for group-average data.

**Supplementary Figure 4.**
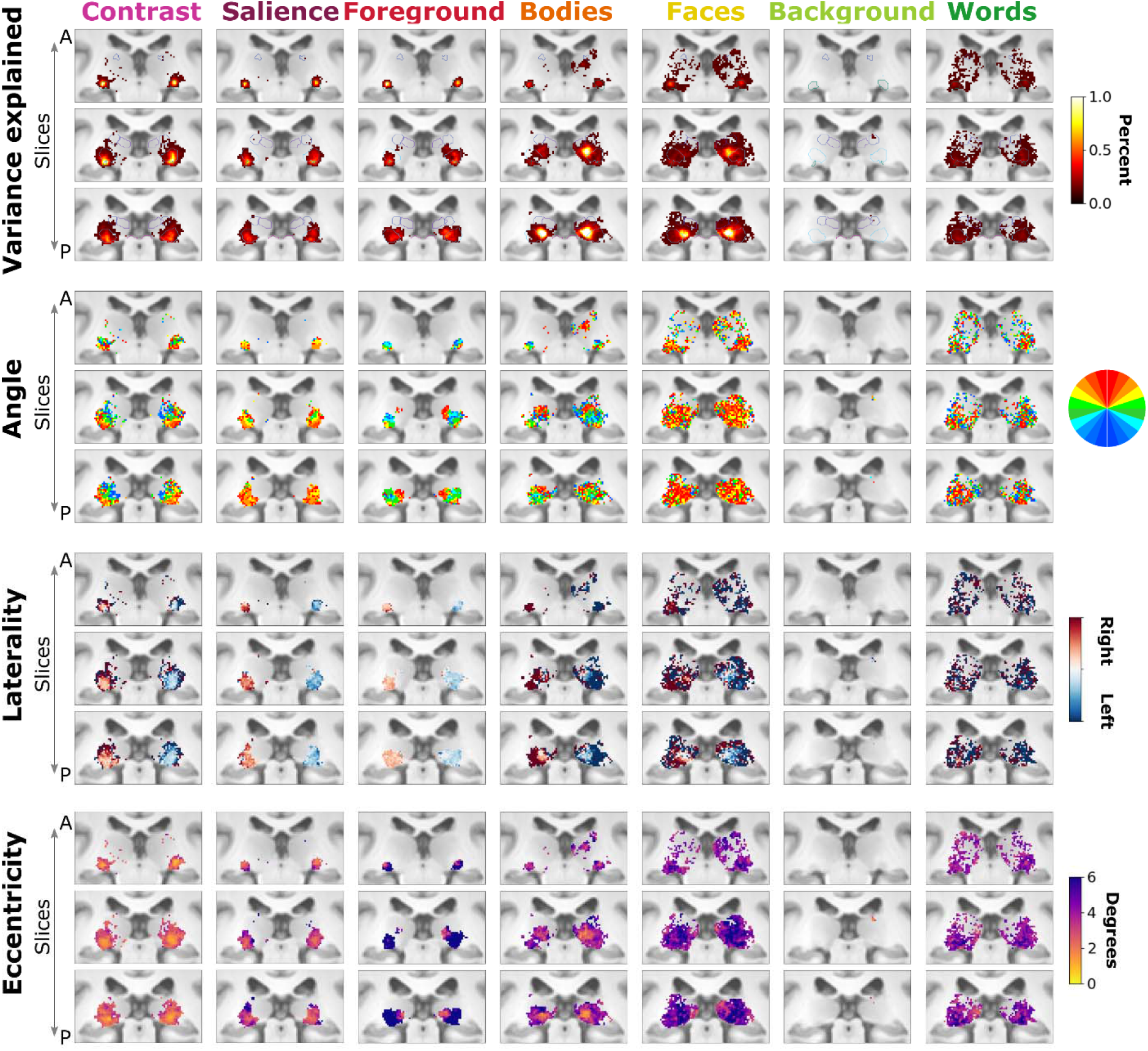
From top to bottom, group-average variance explained, angle, laterality, and eccentricity estimates in three coronals slices (rows) from each tested pRF model (columns) presented as in Figure 3.

**Supplementary Figure 5.**
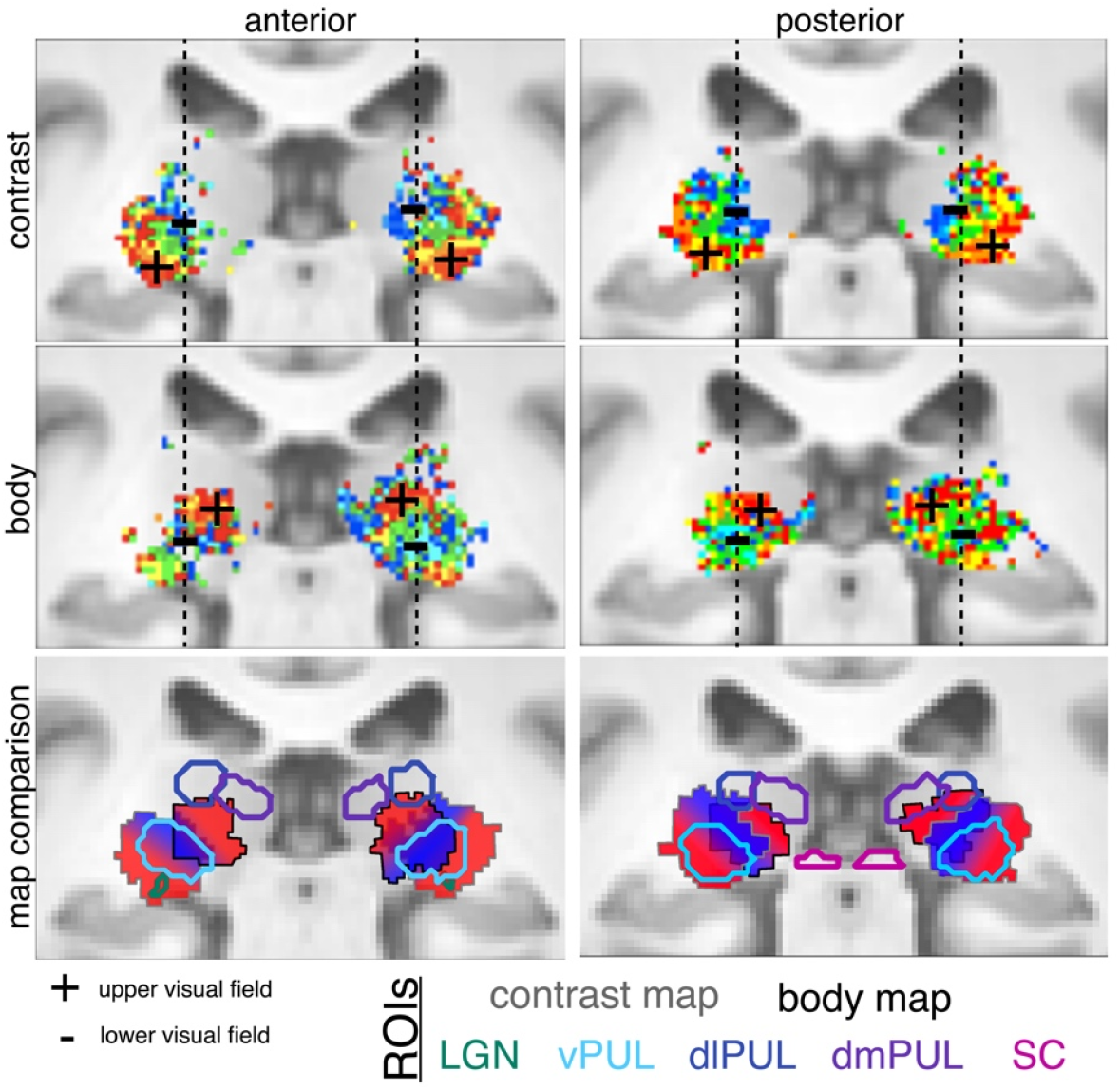
Distinct retinotopic maps for low- and high-level visual features in the medial pulvinar. Polar angle maps derived from the (top) contrast and (middle) body pRF models for two coronal sections within the pulvinar. Upper visual field (UVF) representations (red-yellow-light green colors, indicated by ‘+’) and lower visual field (LVF) representations (blue-light blue-dark green colors, indicated by ‘-’) are shown. A vertical dashed line in each coronal image illustrates the correspondence of the LVF representation between contrast and body maps. (bottom) Schematic of mirror-map organization for low- and high-level visual features. Polar angle organization of lateral and medial ventral pulvinar, derived from contrast and body pRF models respectively, are overlaid in two adjacent coronal sections. Upper and lower visual field (VF) representations are shown as a red-to-blue gradient. The lateral (contrast-derived) and medial (body-derived) pulvinar gradients are inverted and offset relative to each other, with overlapping lower visual field (blue) representations, but distinct upper visual field representations, demonstrating mirror symmetry in the map organization between these pulvinar regions. Colored contours from A are overlaid on maps and show correspondence between previously reported retinotopic organization in the pulvinar (Arcaro et al., 2015) and the contrast map. The body-derived map fell largely outside of these functional ROIs.

